# Investigating the Intradermal Irritation Test of Hydrogel: A Study on Domestic Production

**DOI:** 10.1101/2024.02.19.581046

**Authors:** Kehinde Odelabu, Christianah Racheal

## Abstract

This study employs the intradermal injection method to assess the skin irritation potential of domestically produced medical hydrogel. Healthy white rabbits received intradermal injections of 0.5% sodium chloride solution (control), 0.5% sodium chloride extract, and olive oil extract as test samples. Erythema and edema were observed at 24 and 72 hours post-injection, and the primary irritation index was determined. Results revealed a primary irritation index of 0 for domestically produced hydrogel, indicating no skin irritation response.

## 1. Introduction

The rapid progress of using hydrogels in modern science and technology has significantly propelled the evolution of materials employed in clinical plastic surgery.[1–9] Among these materials, poly(methyl methacrylate) stands out as a noteworthy hydrogel with unique characteristics. This hydrogel is hydrophilic, meaning it has an affinity for water, and possesses the properties of being colorless and transparent, making it a suitable choice for various applications in plastic surgery.[10–12]

Poly(methyl methacrylate), commonly referred to as PMMA, is specifically designed to address tissue depressions both on the surface and internally within the body. Its effectiveness in filling and volumizing areas with depressions or irregularities has led to its widespread use in aesthetic and reconstructive procedures.[13, 14] The hydrophilic nature of PMMA allows it to interact favorably with bodily fluids, contributing to its stability and performance as a filling material.[15]

The synthesis of poly(methyl methacrylate) involves a low-temperature reaction between acrylamide and ammonia.[16] While this process yields a hydrogel with desirable properties, it’s important to note that there might be residual acrylamide monomers present.[17] These residual monomers have the potential to be irritating to the skin, and in some cases, they may trigger toxic reactions that could adversely affect the nerves and muscles.[17] Ensuring the safety of clinical applications requires a comprehensive understanding of these potential risks and the implementation of stringent testing protocols.[18]

To guarantee the safety of using poly(methyl methacrylate) in clinical settings, it becomes imperative to conduct thorough evaluations for potential irritation reactions.[19] Rigorous testing methodologies need to be employed to assess the hydrogel’s safety profile, considering factors such as skin sensitivity, inflammatory responses, and the overall biocompatibility of the material.[20, 21] These evaluations are crucial in determining the hydrogel’s suitability for use in plastic surgery and ensuring that patients are not exposed to unnecessary risks during medical procedures.[22, 23]

As poly(methyl methacrylate) continues to play a significant role in clinical plastic surgery, it is essential to stay abreast of advancements in testing methodologies and safety assessments.[24, 25] This approach ensures that the benefits of using this hydrogel in medical applications are maximized, while potential risks are minimized to guarantee the highest standards of patient safety and well-being.

There are many methods to investigate irritation in animals, including the use of artificial intelligence for simulation and statistics analysis.[26–31] In this article, we applied poly(methyl methacrylate) hydrogels to the animals to observe any resulting irritation.

## 2. Experiment

### 2.1 Materials

Methyl methacrylate monomer, poly(methyl methacrylate) (PMMA), 0.9% sodium chloride injection, Olive oil, New Zealand healthy white rabbits (both genders with weighing not less than 2 kg). The control group consists of 2 animals, and the PMMA sample group consists of 3 animals.

### 2.2 Methods

Control Solution was prepared by mixing 0.06g of methyl methacrylate with 50 ml of 0.5% sodium chloride and sterilized at 121°C for 0.5 hours, following the concentration requirements for residual methyl methacrylate in poly(methyl methacrylate) products. PMMA samples (5g each) were prepared by adding 50ml of 0.5% sodium chloride and 50 ml of olive oil, respectively. The mixtures were sterilized at 121°C for 1 hour, and the supernatant was used as the sample extract within 24 hours. Negative Control Solutions were prepared by adding 20ml each of 0.9% sodium chloride and olive oil, and were separately sterilized at 121°C for 0.5 hours.

### 2.3 Procedures

Control Group: Twenty-four hours before the experiment, 5cm x 5cm areas on both sides of the rabbit spine were shaved. Skin disinfection was performed using 75% ethanol. On one side, 5 points were selected, and 0.2ml of 0.4% sodium chloride was injected at each point. On the other side, 10 points were chosen, and acrylamide control solution was injected. Skin reactions were observed at 24, 48, and 72 hours. PMMA Sample Group: Similar to the control group, but using PMMA sample extract, 0.5% sodium chloride, and olive oil for injections. Skin reactions were observed at the same intervals.

## 3. Results and Discussion

Methyl methacrylate shows noticeable skin irritation reactions at 2-3 injection points when the methyl methacrylate monomer concentration was 2 × 10^−3^ g/ml. PMMA Hydrogel shows no erythema or swelling at injection points for PMMA/0.5% sodium chloride and PMMA/olive oil compared to controls, indicating a low residual methyl methacrylate concentration. The Primary Irritation Index (PII)

**Table 1.** Intradermal reaction grading results for PMMA hydrogels.

| Group | # of injection points | 24 h |  |  |  | 48 h |  |  |  | 72 h |  |  |  |
| --- | --- | --- | --- | --- | --- | --- | --- | --- | --- | --- | --- | --- | --- |
|  |  | 1 | 2 | 3 | 4 | 1 | 2 | 3 | 4 | 1 | 2 | 3 | 4 |
| 0.5% sodium chloride | 10 | - | - | - | - | - | - | - | - | - | - | - | - |
| PMMA/0.5% sodium chloride | 20 | 2** | 2 | - | - | 3 | - | - | - | 2 | 1 | - | - |

\*At each specified time, intradermal reactions (erythema, edema) are graded on a scale of 1 to 4.
\*\*The total number of scores for each grade within the designated.

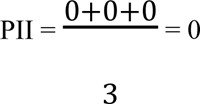

for PMMA, PII was 0, suggesting no potential skin irritation and demonstrating promising application prospects. PMMA/0.5% sodium chloride and PMMA/olive oil showed visible papules protruding beyond the skin surface after injection, indicating good hydrophilicity for PMMA, particularly with 0.5% sodium chloride. Overall, poly(methyl methacrylate) hydrogel exhibits good hydrophilicity and potential for clinical tissue filling.

